# Age-dependent progression of SARS-CoV-2 infection in Syrian hamsters

**DOI:** 10.1101/2020.06.10.144188

**Authors:** Nikolaus Osterrieder, Luca D. Bertzbach, Kristina Dietert, Azza Abdelgawad, Daria Vladimirova, Dusan Kunec, Donata Hoffmann, Martin Beer, Achim D. Gruber, Jakob Trimpert

**Affiliations:** Institut für Virologie, Freie Universität Berlin, Robert-von-Ostertag-Str. 7-13, 14163 Berlin, Germany; Institut für Veterinärpathologie, Freie Universität Berlin, Robert-von-Ostertag-Str. 15, 14163 Berlin, Germany; Institut für Virusdiagnostik, Friedrich-Loeffler-Institut, Südufer 10, 17493 Greifswald - Insel Riems, Germany

**Author notes:** Shared first authorship (Nikolaus Osterrieder, Luca D. Bertzbach). Corresponding author: Jakob Trimpert.

**Keywords:** Coronavirus, *Mesocricetus auratus*, animal model, COVID-19, pneumonia, age-related disease, histopathology, in situ hybridization, cellular tropism, serology

## Abstract

In late 2019, an outbreak of a severe respiratory disease caused by an emerging coronavirus, SARS-CoV-2, resulted in high morbidity and mortality in infected humans^1^. Complete understanding of COVID-19, the multi-faceted disease caused by SARS-CoV-2, requires suitable small animal models, as does the development and evaluation of vaccines and antivirals^2^. Because age-dependent differences of COVID-19 were identified in humans^3^, we compared the course of SARS-CoV-2 infection in young and aged Syrian hamsters. We show that virus replication in the upper and lower respiratory tract was independent of the age of the animals. However, older hamsters exhibited more pronounced and consistent weight loss. *In situ* hybridization in the lungs identified viral RNA in bronchial epithelium, alveolar epithelial cells type I and II, and macrophages. Histopathology revealed clear age-dependent differences, with young hamsters launching earlier and stronger immune cell influx than aged hamsters. The latter developed conspicuous alveolar and perivascular edema, indicating vascular leakage. In contrast, we observed rapid lung recovery at day 14 after infection only in young hamsters. We propose that comparative assessment in young versus aged hamsters of SARS-CoV-2 vaccines and treatments may yield valuable information as this small-animal model appears to mirror age-dependent differences in human patients.

## Main Text

Emerging coronaviruses^4^ have caused serious global public health concerns in the past two decades and cause infections that lead to severe respiratory and partly systemic disease. These include severe acute respiratory syndrome (SARS)-CoV as well as Middle East respiratory syndrome (MERS)-CoV, both of which resulted in high morbidity and mortality in infected humans^5^. Similar to other emerging CoV, the novel SARS-CoV-2 likely arose from an ancestor in bats and amplified in an as yet unknown animal reservoir before making its jump into the human population^6^. SARS-CoV-2 has pushed global health systems to the brink of breakdown. The remarkably fast and unexpected spread of SARS-CoV-2 can be attributed to efficient replication in the upper respiratory tract and robust human-to-human transmission, a characteristic that clearly distinguishes SARS-CoV-2 from SARS- and MERS-CoV. While COVID-19 is primarily a respiratory syndrome, it can induce quite variable clinical signs. It has become evident that differences in type and severity of SARS-CoV-2-induced disease seems to be correlated with the age of patients and is exacerbated by pre-existing conditions^4,7^.

The availability of reliable animal models is of critical importance for pathogenesis studies as well as the development and preclinical evaluation of vaccines and therapeutics. For SARS-CoV-2, the susceptibility of several animal species was predicted by *in silico* analysis based on comparisons of the entry receptor for SARS-CoV and SARS-CoV-2, human angiotensin converting enzyme 2 (hACE2). More specifically, the interaction of the viral spike (S) glycoprotein receptor binding domain with its hACE2 counterpart was examined^8,9^, and in some cases examined *in vivo*^10^. Productive SARS-CoV-2 infection was shown in non-human primates, which developed respiratory disease recapitulating moderate disease as observed in humans^11-14^. Mice are not naturally susceptible to SARS-CoV-2, but mouse-adapted virus strains have been developed and used in BALB/c mice^15,16^. Moreover, transgenic mice expressing hACE2 represent a lethal SARS-CoV-2 infection model resulting in significant weight loss and permitting robust virus replication in the respiratory tract including the lungs^17^. Ferrets have provided valuable data in the case of SARS-CoV^18,19^, and two studies describe the infection of ferrets with SARS-CoV-2 and successful transmission to in-contact animals without clinical signs^20,21^.

First and preliminary studies also focused on the assessment of a Syrian hamster model that had previously been used successfully in SARS and MERS research^18,19,22,23^. It was suggested that hamsters are highly susceptible although they were reported to show no or only moderate respiratory signs and body weight losses. It is important to note, however, that only young male hamsters of 4 to 5 weeks of age were used in these studies^24,25^.

We sought to explore age-related differences in the course of SARS-CoV-2 infection in Syrian hamsters, and to establish a small-animal model that resembles the more severe SARS-CoV-2-infection observed particularly in elderly patients. For our experiments, we used a total of 36 female and male Syrian hamsters (*Mesocricetus auratus*), which were either 6-(n=24) or 32- to 34-week-old (n=12). Hamsters were kept in individually ventilated cages (IVCs), and randomly assigned to three groups: mock (n=12, 6-week-old), young infected (n=12, 6-week-old) and aged infected (n=12, 32- to 34-week-old). Animals were mock-infected with supernatants of cell culture medium taken from uninfected Vero E6 cells or infected with 1×10^5^ plaque-forming units of SARS-CoV-2 München (SARS-CoV-2M; BetaCoV/Germany/BavPat1/2020)^26^. During the 14-day experiment, body temperatures, body weights and clinical signs were recorded daily. Animals were euthanized and sampled at different time points after infection to assess virus titers in various organs and to examine pathological changes in the lungs (Fig. 1 and 2).

**Figure 1:**
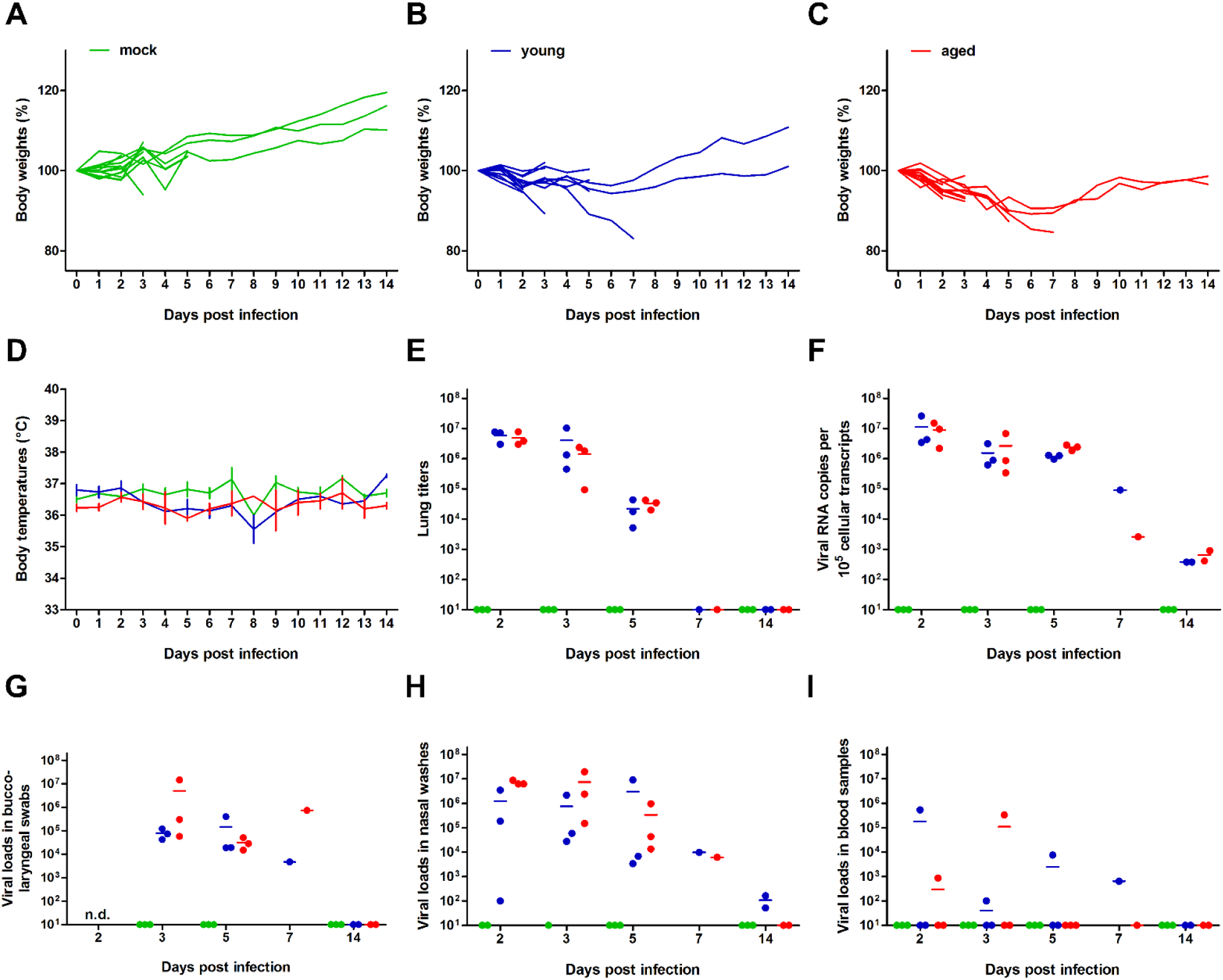
Body weight changes, body temperatures, and viral loads of young versus aged Syrian hamsters infected intranasally with SARS-CoV-2. Individual relative body weights of (**A**) mock-infected, (**B**) young and (**C**) aged hamsters over the course of 14 days after infection are given. (**D**) Temperature changes (as means with SEM). Viral loads were determined from homogenized right cranial lung lobes. (**E**) Virus titers of 25 mg of tissue determined by plaque assay in Vero E6 cells, and (**F**) corresponding virus genome copy numbers as determined by RT-qPCR. Viral loads were also determined by RT-qPCR in (**H**) bucco-laryngeal swabs, (**I**) nasal washes and (**G**) 25 µl of whole blood samples. The color codes represent mock-infected, infected young (6-week-old, blue) and aged hamsters (33- to 35-week-old, red).

**Figure 2:**
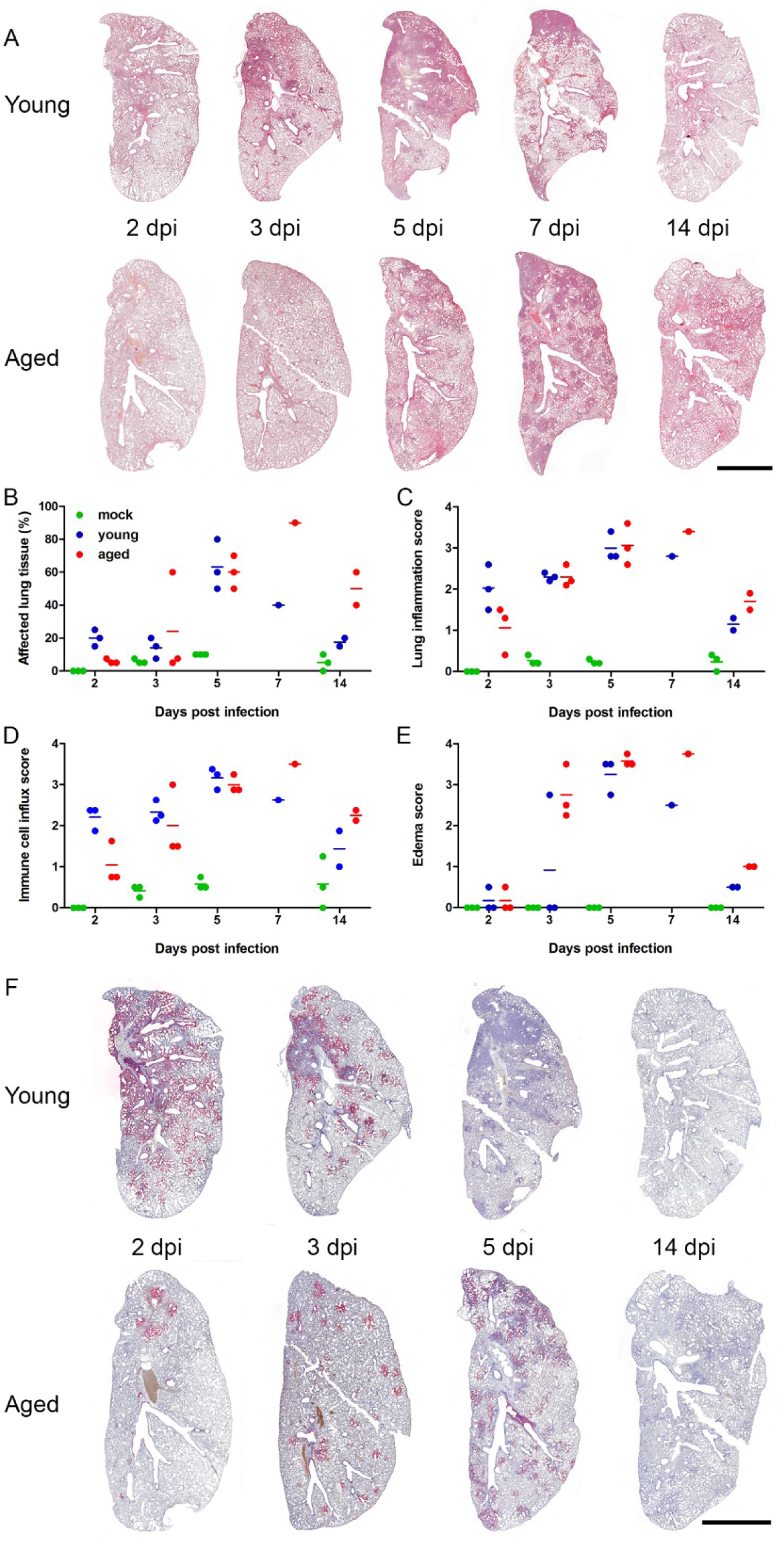
Lung histopathology (A-E) and detection of SARS-CoV-2 RNA (F) at different time points after infection. (**A**) Time-dependent course of pneumonia in young and adult hamster (Bar = 0.5 cm) (**B**) Semiquantitation of lung lesions as assessed by histopathology scores from 0 (absent) to 4 (severe changes) for the following parameters: (**C**) Lung inflammation score taking into account i) severity of pulmonary inflammation; ii) bronchitis (iii) bronchial and alveolar necrosis; iv) hyperplasia of alveolar epithelial cells type II. (**D**) Immune cell influx score taking into account the infiltration of lung tissue with i) neutrophils; ii) macrophages; iii) lymphocytes; iv) perivascular lymphocytic cuffing; and (**E**) edema score including i) alveolar and ii) perivascular edema. (**F**) Time-dependent distribution of SARS-CoV-2 RNA signals in young and adult hamsters as detected by *in situ*-hybridization. (Bar = 0.5 cm)

First, we observed age-dependent SARS-CoV-2-induced body weight losses, with more pronounced weight reductions in aged compared to young hamsters (Fig. 1A-C). Mean body weight losses peaked at 6 to 7 days post-infection (dpi), with partial recovery until 14 dpi in both infected groups. There were no differences in body temperatures between the infected groups or between infected and mock-infected animals (Fig. 1D).

Next, we determined viral titers and SARS-CoV-2 RNA copy numbers in various tissues by RT-qPCR^27,28^, and performed virus titrations from lung homogenates using Vero E6 cells (Fig. 1E-H). Results were similar between age groups and confirmed high viral loads in respiratory samples at early time points after infection, but relatively rapid clearance of infection. It seems important to note that RT-qPCR of blood samples revealed viremia in two male individuals from both age groups with viral RNA copy numbers of >10^5^ at 5 and 7 dpi, respectively (Fig. 1I). In these individuals, we also detected relatively high levels of viral RNA in the spleen, kidneys and duodenum indicating systemic spread of SARS-CoV-2 in some cases (Table S1). To further investigate potential dissemination of infection, we tested the aforementioned organs from all animals sacrificed at 5 dpi and found them to be either negative for SARS-CoV-2 RNA or to contain only low levels of viral RNA (Tab. S1).

Viral loads in bucco-laryngeal swabs and nasal washes appeared to be a reliable surrogate of viral loads in the lungs in both age groups, as the average loads ranged between 10^4^ and 10^7^ copies, respectively, at early times after infection, indicating that these sampling techniques can be used to monitor SARS-CoV-2 replication in Syrian hamsters (Fig. 1G-H).

At 14 dpi, hamsters had mounted a robust humoral immune response as evidenced by relatively high titers of neutralizing antibodies. It is worth noting that antibody titers were higher in young when compared to aged hamsters (Table S1).

Histopathology revealed clear age-dependent differences, with young hamsters launching an earlier and stronger immune cell influx into the lungs associated with a faster recovery than their aged counterparts (Fig. 2A-E). At 2 dpi, young hamsters developed a marked necro-suppurative bronchointerstitial pneumonia with strong alveolar and interstitial influx of neutrophils and macrophages as well as perivascular lymphocytic cuffing, which was much milder or absent in the aged group (Fig. S1A, right panel). In contrast, only aged animals developed pronounced alveolar and perivascular edema indicating vascular leakage at 3 dpi (Fig. S2C). A more diffuse, severe bronchointerstitial pneumonia was similarly present in both groups at 5 dpi with onset of tissue regeneration, including hyperplasia of bronchial epithelium (Fig. S1C, arrowhead) and type II alveolar epithelial cells.

Only at that time point, the arterial and venous endothelium of animals in both groups was swollen and vacuolated, with necrotic endothelial cells separated from the underlying basement membrane by the presence of subendothelial lymphocytes and neutrophils (Fig. S2C, left), consistent with what has been described as endothelialitis in human SARS-CoV-2 infection^29^. At 7 dpi, recovery as indicated by marked hyperplasia of bronchial epithelial cells and type II alveolar epithelial cells were seen in both groups (Fig. S2A). Interestingly, lung tissues had almost recovered in young hamsters at day 14, while the aged animals still had persistent tissue damage and active inflammation (Fig. S2B).

From 2 dpi onwards, SARS-CoV-2 RNA was detected by *in situ* hybridization in bronchial epithelial cells, debris in the bronchial lumen, alveolar epithelial cells type I and type II as well as macrophages in both groups, again with clear age-dependent differences over time (Fig. 2F). Young animals had high amounts of viral RNA in numerous bronchial epithelial cells and within the bronchial lumen that was accompanied by marked spreading through the lung parenchyma on 2 and 3 dpi. In contrast, aged animals, had less virus RNA present in the bronchi. We detected only a scattered pattern of infected bronchial epithelial cells and sporadic areas of parenchymal infection at 2 and 3 dpi. At 5 dpi, viral RNA was undetectable in the bronchi of young hamsters, and only small infected areas containing low levels of RNA with a patchy distribution were detected. It is noteworthy that aged animals, at the same time after infection, had increased numbers of infected areas with a similarly patchy distribution throughout the lungs as well as copious amounts of viral RNA associated with cellular debris in the bronchial lumen. Using this technique, no viral RNA was detected at 14 dpi in either group.

In summary, our study examined the suitability of a small animal model to study SARS-CoV-2 infections. Intranasal infection of Syrian hamsters resulted in weight loss and robust virus replication in the upper and lower respiratory tract. We further demonstrate that aged 32-to 34-week-old hamsters experienced higher and more consistent weight loss after intranasal infection, while body temperatures and virus replication in upper airways and lungs were similar between both age groups. Furthermore, we show that, using *in situ* hybridization, viral RNA was detectable in bronchial epithelial cells, type I and type II alveolar epithelial cells, and macrophages. All these cell types are potential targets of SARS-CoV-2 in human lung tissue; hence, infection of hamsters of different ages seems to closely reflect what has been reported for human patients^25,30^. In contrast to SARS-CoV-2 titers, histopathological changes differed markedly between young and aged Syrian hamsters over time: younger animals launched more severe reactions at early time points after infection, while lesions and inflammation in the lungs became more pronounced and widespread at later time points in the elderly.

Based on the data presented here, we propose that comparative preclinical assessments of SARS-CoV-2 vaccines and other treatment options in young versus aged hamsters may yield valuable and relevant results as this small animal model appears to mimic age-dependent differences in humans. The development of a substantial humoral immune response emphasizes that hamsters are likely suitable for vaccination trials. Our observations also confirm that body weight loss appears to be the only robust clinical parameter in SARS-CoV-2 infection of Syrian hamsters. This makes the difference in body weight loss between age groups with more consistent losses in aged hamsters only more important as it provides an objective way to judge clinical efficacy of antiviral therapy or vaccination.

## Acknowledgements

The authors acknowledge the excellent technical assistance by Ann Reum, Annett Neubert, and Simon Dökel. We would like to thank Carfil Inc. for their generous support of our animal husbandry. This research was supported by COVID-19 grants from Freie Universität Berlin and Berlin University Alliance to NO as well as Deutsche Forschungsgemeinschaft grant SFB-TR84 awarded to ADG.

## Author contributions

Conceptualization: NO, LDB, ADG and JT

Investigation: NO, LDB, KD, AA, DV, DH, DK and JT

Writing: NO, LDB, KD, MB, ADG, and JT

Editing: all authors had the opportunity to comment on the draft manuscript.

