## Supplemental Information for "Age-dependent progression of SARS-CoV-2 infection in Syrian hamsters"

This file includes:
Materials and Methods
Figures S1 and S2
Tables S1 and S2
SI References

### Materials and Methods

#### Ethics statement

We conducted all animal work in compliance with relevant national and international guidelines for care and humane use of animals. The animal use protocol for the experiments reported here was approved by the Landesamt für Gesundheit und Soziales in Berlin, Germany (approval number 0086/20).

#### Viruses and cells

Virus stocks were prepared from a previously published SARS-CoV-2 isolate (BetaCoV/Germany/BavPat1/2020)[1], which was kindly provided by Drs. Daniela Niemeyer und Christian Drosten, Charité Berlin, Germany. The isolate, referred to as SARS-CoV-2 München (SARS-CoV-2M)[2], was handled under the appropriate safety precautions in a BSL-3 facility (Freie Universität Berlin, Institut für Virologie) and propagated on Vero E6 cells (ATCC CRL-1586) in minimal essential medium (MEM; PAN Biotech, Aidenbach, Germany) supplemented with 10% fetal bovine serum (PAN Biotech), 100 IU/ml penicillin G and 100 µg/ml streptomycin (Carl Roth, Karlsruhe, Germany).

#### Animal Experiments

Thirty-six 6- or 32-to-34-week-old female and male Syrian hamsters (*Mesocricetus auratus*; breed RjHan:AURA, Janvier Labs, Saint-Berthevin, France) were kept in individually ventilated cages (IVCs; Tecniplast, Buguggiate, Italy) in an approved BSL-3 facility. IVCs were equipped with enrichment (Carfil, Oud-Turnhout, Belgium). All animals had unrestricted access to food and water and were allowed to acclimate to the conditions for seven days prior to infection. Cage temperatures and relative humidities were recorded daily and ranged from 22-24°C and 40-55%, respectively.

Animal experiments

The 6-week-old hamsters were randomly distributed into two groups: mock (n=12, 6-week-old), young infected (n=12, 6-week-old). The third group represents the aged infected hamsters (n=12, 32-34-week-old). IPTT-300 transponders (BioMedic Data Systems, Seaford, DE, USA) were subcutaneously implanted into all hamsters 2 days prior to infection to allow identification and monitoring of body temperatures. Animals were mock-infected with 60 µl medium from uninfected Vero E6 cells or infected with $1X10^5$ pfu SARS-CoV-2M in 60 µl by intranasal instillation. For transponder implantation, hamsters were sedated with butorphanol (2.5 mg/kg; CP-Pharma, Burgdorf, Germany) and midazolam (2 mg/kg; Braun, Melsungen, Germany). For infections, hamsters were sedated with ketamine (25 mg/kg; Serumwerk Bernburg, Bernburg, Germany) and midazolam (2 mg/kg; Braun).

On 2, 3 and 5 days post-infection (dpi), three hamsters of each group were euthanized by exsanguination under medetomidine (0.15 mg/kg; Pharma-Partner, Hamburg, Germany), midazolam (2 mg/kg) and butorphanol (2.5 mg/kg) anesthesia[3]. Blood, nasal washes, bucco-laryngeal swabs, lungs (left and right), kidneys, spleens, duodenums and blood sera were collected for (histo)pathological examinations and/or virus titrations, RT-qPCR and serological examination. During the 14-day experiment, body temperatures, body weights and clinical signs of all animals were monitored twice daily. Animals that with a body weight loss of more than 10% weight over a 72 h period were euthanized in compliance with the animal use protocol. Such humane termination applies to the two hamsters euthanized 7 dpi.

Histopathological examination

For histopathology and *in situ* hybridization (ISH) the left lung lobe was carefully removed, immersion-fixed in formalin, pH 7.0, for 48 h, embedded in paraffin, and cut in 2 µm sections. For histopathology, slides were stained with hematoxylin and eosin (HE) after dewaxing in xylene and rehydration in decreasing ethanol concentrations. Lung sections were microscopically evaluated in a blinded fashion by a board-certified veterinary pathologist to assess character and severity of pathologic lesions using lung-specific inflammation scoring parameters as described for other lung infection models before[4]. Three different scores were used that included the following parameters: (1) lung inflammation score including severity of (i) interstitial pneumonia (ii) bronchitis, (iii) epithelial necrosis of bronchi and alveoli, and (iv) hyperplasia of type II-alveolar epithelial cells; (2) immune cell infiltration score taking into account the presence of (i) neutrophils; (ii) macrophages, and (iii) lymphocytes in the lungs as well as (iv) perivascular lymphocytic cuffing; and (3) edema score including (i) alveolar edema and (ii) perivascular edema.

ISH was performed as reported previously[5] using the ViewRNA™ ISH Tissue Assay Kit (Invitrogen by Thermo Fisher Scientific, Darmstadt, Germany) following the manufacturer's instructions with minor adjustments. Probes for the detection of N gene RNA of SARS-CoV-2 (NCBI database NC_045512.2, nucleotides 28,274 to 9,533, assay ID: VPNKRHM) and the mouse housekeeping gene eukaryotic translation elongation factor-1α (EF1a; assay ID: VB1-14428-VT, Affymetrix, Inc., Santa Clara, CA, USA), which that shares 95% sequence identity with the Syrian hamster orthologue, were designed. Lung sections (2 µm thickness) on adhesive glass slides were dewaxed in xylol and dehydrated in ethanol. Tissues were incubated at 95°C for 10 min with subsequent protease digestion for 20 min. Sections were fixed with 4% paraformaldehyde in PBS (Alfa Aesar, Thermo Fisher, Kandel, Germany) and hybridized with the probes. Amplifier and label probe hybridizations were performed according to the manufacturer's instructions using fast red as the chromogen, followed by counterstaining with hematoxylin for 45 s, washing in tap water for 5 min, and mounting with Roti®-Mount Fluor-Care DAPI (4, 6-diaminidino-2-phenylindole; Carl Roth). For negative and morphologically intact controls, lungs from uninfected hamsters of each group (n=4) were included. In addition, an irrelevant probe for the detection of pneumolysin was used as a negative control for unspecific reactions. HE-stained and ISH slides were analyzed and images taken using an Olympus BX41 microscope with a DP80 Microscope Digital Camera and the cellSens™ Imaging Software, Version 1.18 (Olympus Corporation, Münster, Germany). For the display of overviews of whole lung lobe sections, slides were automatically digitized using the Aperio CS2 slide scanner (Leica Biosystems Imaging Inc., Vista, CA, USA), and image files were generated using the Image Scope Software (Leica Biosystems Imaging Inc.).

Virus titrations

For assessment of virus titers from 25 mg of lung tissue, tissue homogenates were serially diluted and plated on Vero E6 cells in 12-well cell culture plates (Sarstedt, Nümbrecht, Germany). At 3 dpi, cells were fixed in 4% formalin, stained with 0.1% crystal violet (in 25% methanol), and plaques were counted.

RNA extractions and quantitative RT-PCR

RNA was extracted from nasal washes and tracheal swabs with the RTP DNA/RNA Virus Mini Kit (Stratec, Birkenfeld, Germany) according to the manufacturer's instructions. The innuPREP Virus DNA/RNA Kit (Analytic Jena, Jena, Germany) was used for RNA extractions from tissue samples. Viral RNA was quantified using a one-step RT qPCR reaction with the NEB Luna Universal Probe One-Step RT-qPCR (New England Biolabs, Ipswitch, MA, USA) and the 2019-nCoV RT-qPCR primers and probe (E_Sarbeco)[6] on a StepOnePlus RealTime PCR System (Thermo Fisher Scientific, Waltham, MA, USA) according to the manufacturer's instructions. Viral RNA copies were then normalized to cellular RPL18 as previously described[7]. All primers and probes are listed in Table S2.

Standard curves for absolute quantification were generated from serial dilutions of SARS-CoV-2 RNA obtained from a full-length virus genome cloned as a bacterial artificial chromosome and propagated in *E. coli* or from serial dilutions of the purified hamster RPL-18 PCR product. The latter was generated by PCR using RPL-18 qPCR-primers (Table S2) and a cDNA template obtained from hamster lung tissue. In lung tissue, viral RNA copies were calculated per $10^5$ hamster RPL-18 transcripts.

Serology

For serum neutralization assays, 50 µl of medium containing $10^{3.3}$ tissue culture infectious doses $_{50}$ ($TCID_{50}$) of SARS-CoV-2M were mixed with 50 µl of diluted serum. Each sample was tested in triplicate. After 1 h incubation at 37°C, the mixture was transferred to confluent Vero E6 cells in a 96-well plate (Sarstedt, Nümbrecht, Germany). Viral replication was assessed after 3 days by the detection of cytopathic effects.

Statistical analyses

Statistical analyses were performed using Graph-Pad Prism v8 (GraphPad Software Inc., San Diego, CA, USA). Statistical details of all analyzed experiments can be found in the respective figure legend.

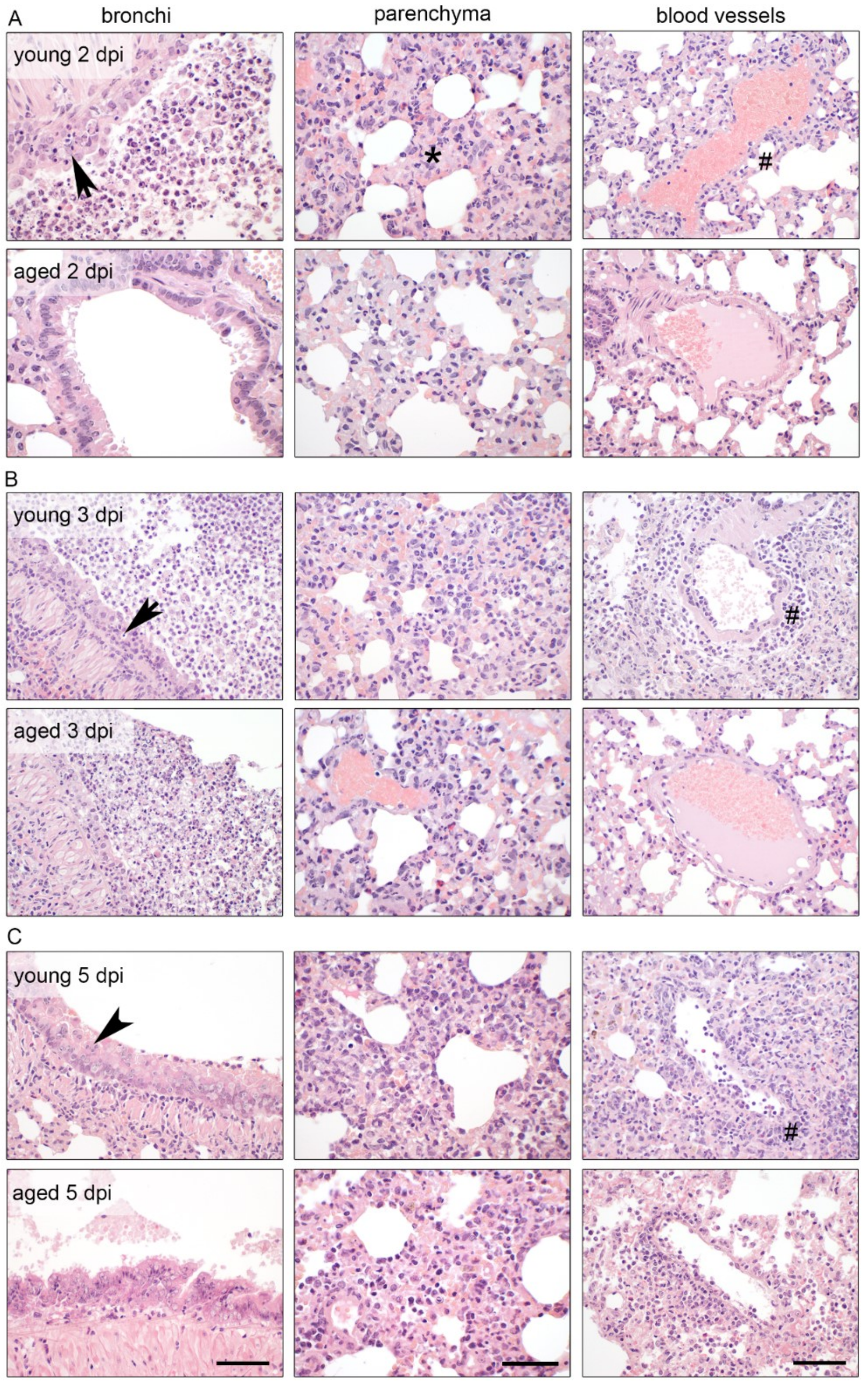


**Figure S1: Histopathological comparison at early time points after SARS-COV-2 infection.** (**A**) At 2 dpi, young animals developed marked necrotizing (left, arrow) and suppurative bronchitis (left) with cell debris and exudates filling the bronchial lumen, widely expanded interstitial tissue (center, asterisk) with infiltration of macrophages, neutrophils and lymphocytes and onset of perivascular lymphocytic cuffing (right, hash), which was milder or absent in aged hamsters. (**B**) At 3 dpi, necro-suppurative bronchitis (left, arrow) as well as interstitial pneumonia accelerated in both groups, while perivascular lymphocytic cuffing present only in young animals (right, hash). (**C**) At 5 dpi, hyperplasia of bronchial epithelia (left, arrowhead), interstitial pneumonia, and perivascular lymphocytic cuffing (right, hash) were identical in young and aged animals. Bars: left panels, 100 µm; center panels, 50 µm; right panels, 100 µm.

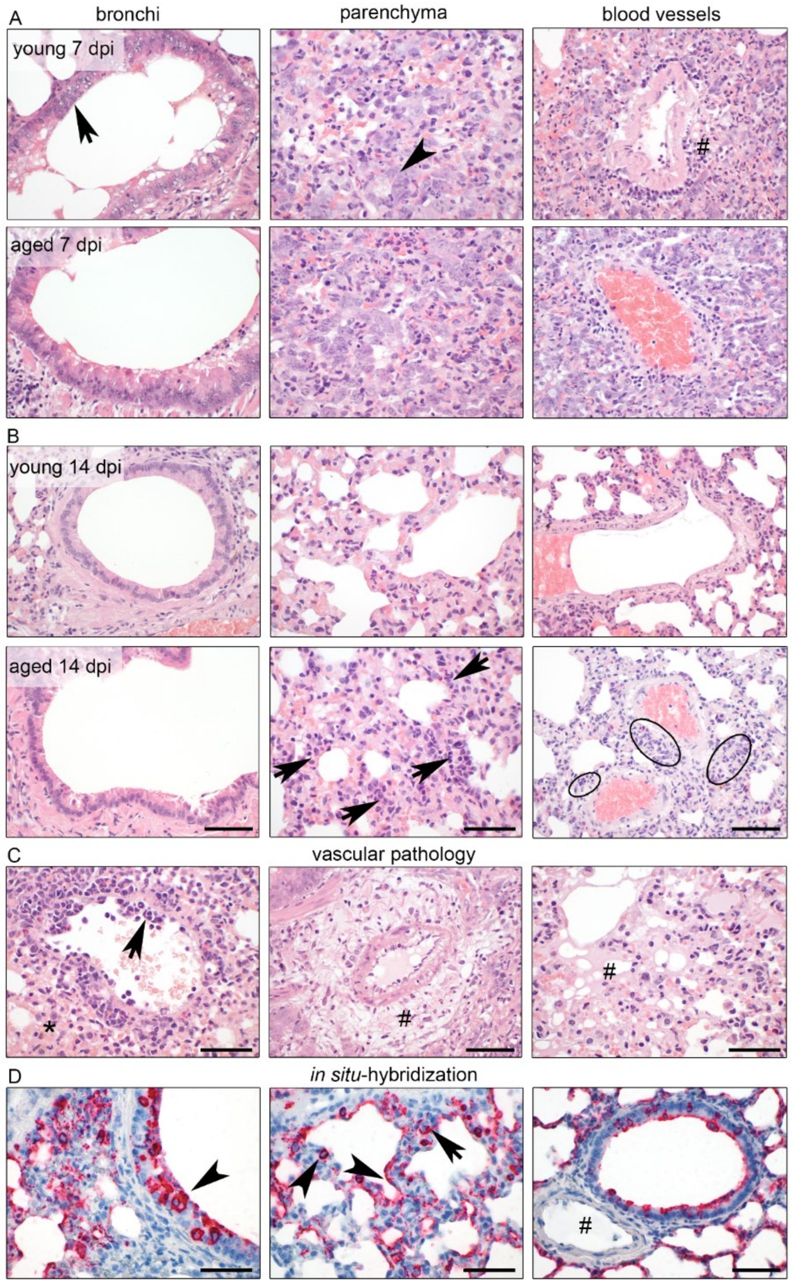


**Figure S2: Histopathological comparison at late time points including vascular pathology and detection of SARS-COV-2 RNA.** (**A**) At 7 dpi, dominant regeneration of bronchial epithelial cells (left, arrow) and type II alveolar epithelial cells (center, arrowhead) as well as prominent perivascular lymphocytic cuffing (right, hash) were present in both groups. (**B**) At 14 dpi, lungs of young animals showed only minimal mononuclear cell infiltration, restored tissue structures and largely resolved inflammation, while lungs of aged hamsters still had persistent tissue damage and active inflammation (center, arrow, right, oval). (**C**) Vascular pathology included endothelialitis (left, arrow), alveolar hemorrhage (left, asterisk), perivascular edema (center, hash) and alveolar edema (right, hash). (**D**) Viral RNA was detected by *in situ* hybridization in bronchial epithelial cells (left, arrowhead), type I and II alveolar epithelial cells (center, arrowhead), and macrophages (center, arrow). Endothelial cells (right, hash) were not infected at the any of the time points in any of the lungs. Bars (A, B): left panels, 100 µm; center panels, 50 µm; right panels, 100 µm. Bars (C): left and right, 50 µm; center, 100 µm. Bars (D): left and center, 50 µm; right, 100 µm.

**Table S1:** Virus titers and RNA copies in lung homogenates (25 mg – RNA copies per $10^5$ cellular transcripts), nasal washes, bucco-pharyngeal swabs, blood samples (25 µl), kidneys (25 mg), spleens (25 mg) and duodenums (25 mg), as well as serum neutralizing antibodies of each hamster.

| # | group | sex | sample date | age | lung titers | RNA copies (lung) | RNA copies (wash) | RNA copies (swab) | RNA copies (blood) | RNA copies (kidney) | RNA copies (spleen) | RNA copies (duodenum) | Neutralizing antibody titer |
|---|---|---|---|---|---|---|---|---|---|---|---|---|---|
| 12 | uninf. | ♀ | 2 dpi | young | 0.00 | 0.00 | 0.00 | n.d. | 0.00 | n.d. | n.d. | n.d. | n.d. |
| 19 | uninf. | ♂ | 2 dpi | young | 0.00 | 0.00 | 0.00 | n.d. | 0.00 | n.d. | n.d. | n.d. | n.d. |
| 31 | uninf. | ♀ | 2 dpi | young | 0.00 | 0.00 | 0.00 | n.d. | 0.00 | n.d. | n.d. | n.d. | n.d. |
| 10 | uninf. | ♀ | 3 dpi | young | 0.00 | 0.00 | 0.00 | 0.00 | 0.00 | n.d. | n.d. | n.d. | n.d. |
| 13 | uninf. | ♀ | 3 dpi | young | 0.00 | 0.00 | 0.00 | 0.00 | 0.00 | n.d. | n.d. | n.d. | n.d. |
| 27 | uninf. | ♂ | 3 dpi | young | 0.00 | 0.00 | 0.00 | 0.00 | 0.00 | n.d. | n.d. | n.d. | n.d. |
| 3 | uninf. | ♀ | 5 dpi | young | 0.00 | 0.00 | 0.00 | 0.00 | 0.00 | 0.00 | 0.00 | 0.00 | n.d. |
| 29 | uninf. | ♂ | 5 dpi | young | 0.00 | 0.00 | 0.00 | 0.00 | 0.00 | 0.00 | 0.00 | 0.00 | n.d. |
| 33 | uninf. | ♂ | 5 dpi | young | 0.00 | 0.00 | 0.00 | 0.00 | 0.00 | 0.00 | 0.00 | 0.00 | n.d. |
| 8 | uninf. | ♀ | 14 dpi | young | 0.00 | 0.00 | 0.00 | 0.00 | 0.00 | n.d. | n.d. | n.d. | <1:8 |
| 11 | uninf. | ♀ | 14 dpi | young | 0.00 | 0.00 | 0.00 | 0.00 | 0.00 | n.d. | n.d. | n.d. | <1:8 |
| 34 | uninf. | ♀ | 14 dpi | young | 0.00 | 0.00 | 0.00 | 0.00 | 0.00 | n.d. | n.d. | n.d. | <1:8 |
| 5 | inf. | ♂ | 2 dpi | young | $3.00x10^6$ | $3.46x10^6$ | $3.45x10^6$ | n.d. | $5.34x10^5$ | $5.89x10^3$ | 0.00 | $3.21x10^3$ | n.d. |
| 15 | inf. | ♂ | 2 dpi | young | $7.60x10^6$ | $4.30x10^6$ | $1.87x10^5$ | n.d. | 0.00 | n.d. | n.d. | n.d. | n.d. |
| 35 | inf. | ♂ | 2 dpi | young | $7.20x10^6$ | $2.60x10^7$ | $1.01x10^2$ | n.d. | 0.00 | n.d. | n.d. | n.d. | n.d. |
| 2 | inf. | ♀ | 3 dpi | young | $1.34x10^6$ | $6.20x10^5$ | $2.74x10^4$ | $7.36x10^4$ | $1.01x10^2$ | n.d. | n.d. | n.d. | n.d. |
| 9 | inf. | ♂ | 3 dpi | young | $1.03x10^7$ | $3.14x10^6$ | $2.17x10^6$ | $4.26x10^4$ | 0.00 | n.d. | n.d. | n.d. | n.d. |
| 26 | inf. | ♀ | 3 dpi | young | $4.49x10^5$ | $8.90x10^5$ | $5.89x10^4$ | $1.21x10^5$ | 0.00 | n.d. | n.d. | n.d. | n.d. |
| 1 | inf. | ♂ | 5 dpi | young | $4.33x10^4$ | $1.29x10^6$ | $6.80x10^3$ | $1.88x10^4$ | $7.56x10^3$ | n.d. | n.d. | n.d. | n.d. |
| 24 | inf. | ♀ | 5 dpi | young | $5.15x10^3$ | $9.56x10^5$ | $3.39x10^3$ | $4.00x10^5$ | 0.00 | 0.00 | 0.00 | 0.00 | n.d. |
| 36 | inf. | ♀ | 5 dpi | young | $1.79x10^4$ | $1.26x10^6$ | $9.07x10^6$ | $1.94x10^4$ | 0.00 | 0.00 | 0.00 | 0.00 | n.d. |
| 25 | inf. | ♂ | 7 dpi | young | 0.00 | $9.18x10^4$ | $9.72x10^3$ | $4.73x10^3$ | $6.43x10^2$ | n.d. | n.d. | n.d. | n.d. |
| 14 | inf. | ♂ | 14 dpi | young | 0.00 | $3.80x10^2$ | $1.64x10^2$ | 0.00 | 0.00 | n.d. | n.d. | n.d. | 1:512 |
| 30 | inf. | ♂ | 14 dpi | young | 0.00 | $3.76x10^2$ | $5.26x10^1$ | 0.00 | 0.00 | n.d. | n.d. | n.d. | 1:203.2 |

| | | | | | | | | | | | | | |
|---|---|---|---|---|---|---|---|---|---|---|---|---|---|
| **18** | inf. | ♂ | 2 dpi | aged | $7.80x10^6$ | $2.20x10^6$ | $6.24x10^6$ | n.d. | 0.00 | n.d. | n.d. | n.d. | n.d. |
| **20** | inf. | ♂ | 2 dpi | aged | $3.00x10^6$ | $1.51x10^7$ | $6.20x10^6$ | n.d. | 0.00 | n.d. | n.d. | n.d. | n.d. |
| **32** | inf. | ♂ | 2 dpi | aged | $3.80x10^6$ | $9.52x10^6$ | $8.84x10^6$ | n.d. | $8.71x10^2$ | n.d. | n.d. | n.d. | n.d. |
| **7** | inf. | ♀ | 3 dpi | aged | $9.42x10^4$ | $6.78x10^6$ | $1.51x10^5$ | $5.81x10^4$ | 0.00 | n.d. | n.d. | n.d. | n.d. |
| **17** | inf. | ♂ | 3 dpi | aged | $2.37x10^6$ | $8.72x10^5$ | $1.96x10^7$ | $3.04x10^5$ | $3.33x10^5$ | $3.76x10^3$ | $3.11x10^3$ | $1.15x10^4$ | n.d. |
| **21** | inf. | ♂ | 3 dpi | aged | $1.80x10^6$ | $3.38x10^5$ | $2.39x10^6$ | $1.46x10^7$ | 0.00 | n.d. | n.d. | n.d. | n.d. |
| **6** | inf. | ♀ | 5 dpi | aged | $2.00x10^4$ | $1.83x10^6$ | $1.35x10^4$ | $1.51x10^4$ | 0.00 | n.d. | n.d. | n.d. | n.d. |
| **23** | inf. | ♀ | 5 dpi | aged | $3.45x10^4$ | $2.84x10^6$ | $4.30x10^4$ | $2.86x10^4$ | 0.00 | 0.00 | $9.27x10^1$ | $6.95x10^2$ | n.d. |
| **28** | inf. | ♂ | 5 dpi | aged | $4.25x10^4$ | $2.44x10^6$ | $9.55x10^5$ | $5.09x10^4$ | 0.00 | $2.56x10^2$ | 0.00 | $2.93x10^2$ | n.d. |
| **16** | inf. | ♀ | 7 dpi | aged | 0.00 | $2.58x10^3$ | $6.19x10^3$ | $7.45x10^5$ | 0.00 | n.d. | n.d. | n.d. | n.d. |
| **4** | inf. | ♀ | 14 dpi | aged | 0.00 | $4.08x10^2$ | 0.00 | 0.00 | 0.00 | n.d. | n.d. | n.d. | 1:128 |
| **22** | inf. | ♀ | 14 dpi | aged | 0.00 | $8.98x10^2$ | 0.00 | 0.00 | 0.00 | n.d. | n.d. | n.d. | 1:80.6 |

Table S2: Oligonucleotides used in this study.

| **Primer/probe** | **Sequence 5'–3'** |
|---|---|
| SARS-CoV-2 forward | ACAGGTACGTTAATAGTTAATAGCGT |
| SARS-CoV-2 reverse | ATATTGCAGCAGTACGCACACA |
| SARS-CoV-2 probe | FAM-ACACTAGCCATCCTTACTGCGCTTCG-BHQ |
| RPL-18 forward | GTTTATGAGTCGCACTAACCG |
| RPL-18 reverse | TGTTCTCTCGGCCAGGAA |
| RPL-18 probe | FAM-TCTGTCCCTGTCCCGGATGATC-BHQ |
